# Dying cell-released exosomal CXCL1 promotes breast cancer metastasis by activating TAM/PD-L1 signaling

**DOI:** 10.1101/2023.02.23.529643

**Authors:** Shengqi Wang, Jing Li, Neng Wang, Bowen Yang, Yifeng Zheng, Xuan Wang, Juping Zhang, Bo Pan, Zhiyu Wang

## Abstract

**Background:** Emerging evidence suggests that dying cell-released signals may induce cancer progression and metastasis by modulating the surrounding microenvironment. However, the underlying molecular mechanisms and targeting strategies are yet to be explored.

**Methods:** Apoptotic breast cancer cells induced by paclitaxel treatment were sorted and their released exosomes (exo-dead) were isolated from the cell supernatants. Chemokine array analysis was conducted to identify the crucial molecules in exo-dead. Zebrafish and mouse xenograft models were used to investigate the effect of exo-dead on breast cancer progression *in vivo*. Multiple molecular biological experiments were conducted to determine the underlying mechanisms of exo-dead in promoting breast cancer, as well as its intervention values.

**Results:** It was demonstrated that exo-dead were phagocytized by macrophages and induced breast cancer metastasis by promoting the infiltration of immunosuppressive PD-L1^+^ TAMs. Chemokine array identified CXCL1 as a crucial component in exo-dead to activate TAM/PD-L1 signaling. Exosomal CXCL1 knockdown or macrophage depletion significantly inhibited exo-dead-induced breast cancer growth and metastasis. Mechanistic investigations revealed that CXCL1^exo-dead^ enhanced TAM/PD-L1 signaling by transcriptionally activating EED-mediated PD-L1 promoter activity. More importantly, TPCA-1 (2-[(aminocarbonyl) amino]-5-(4-fluorophenyl)-3-thiophenecarboxamide) was screened as a promising inhibitor targeting exosomal CXCL1 signals to enhance paclitaxel chemosensitivity and limit breast cancer metastasis without noticeable toxicities.

**Conclusions:** Our results highlight CXCL1^exo-dead^ as a novel dying cell-released signal and provide TPCA-1 as a targeting candidate to improve breast cancer prognosis.

## 1. Introduction

Breast cancer is the most commonly diagnosed malignancy and the leading reason for cancer-associated mortality among women worldwide ^1^. Breast cancer alone resulted in 2.3 million new cancer cases and 685,000 deaths in 2020 worldwide, accounting for 24.5% of new cancer cases and 15.5% of cancer deaths among women ^1^. Despite the significant advances in therapeutic strategies in recent decades, cytotoxic chemotherapy represents the cornerstone treatment for patients with breast cancer ^2, 3^, especially the advanced or metastatic cases ^4^. It is estimated that the standard chemotherapy regimen could reduce the 10-year mortality of breast cancer by one-third ^5^. However, the clinical efficacy of chemotherapy in patients with breast cancer is limited by the low response rate in some cases. Indeed, the response rate of taxanes for metastatic breast cancer ranges from 30–70% ^6^. More importantly, chemotherapy often leads to secondary multidrug resistance, resulting in recurrence or metastasis. Notably, emerging studies have suggested that chemotherapy promotes breast cancer immune escape *via* stress-related machinery ^7–10^. Therefore, it is important to further investigate the influence of chemotherapy on the biological behaviour of breast cancer cells and the underlying molecular mechanisms.

Cell death is a biological process that is fundamental to maintaining organismal homeostasis and defending against infection and cancer. It is estimated that more than 100 billion cells die and are renewed in the human body every day ^11^. Dying cells are not simply inert cells waiting for removal, but instead, can release intracellular components as “goodbye” signals that actively modulate cellular fate in surrounding tissues ^12^. Dying cell-released signals are multitudinous, and include metabolic molecules, cytokines, chemokines, proteins, nucleic acids, ion signals and various vesicles. Increasing evidence has indicated that dying cell-released signals are closely involved in cancer metabolism, angiogenesis, drug-resistance, metastasis, and tumor immunity. Dying cell-released signals mainly act as survival signals that promote the survival of surrounding cancer cells. Indeed, dying pancreatic cancer cells exhibit increased release of miR-194-5p, which promotes tumor survival and metastasis by activating the proliferation of residual tumor repopulating cells ^13^. More importantly, dying cell-released signals often act as important messengers to regulate cancer immunogenicity, inflammatory cell infiltration, and immune response activity in the tumor microenvironment (TME). Dying cancer cells release numerous immunostimulatory damage-associated pattern molecules (DAMPs) as “find me” and “eat me” signals, which recruit and activate dendritic cells (DCs) or macrophages to trigger immune responses ^14^. Dying cancer cells can also upregulate the expression and release of tumor-associated antigens and pro-inflammatory cytokines, leading to enhanced immune responses and improved immunotherapy outcomes ^15, 16^. Increasing studies have indicated that chemotherapy might facilitate cancer immune escape and metastasis. However, precise molecular mechanisms remain to be explored. Chemotherapy inevitably results in massive amounts of tumor cell death, leading to the release of multiple dying signals. Therefore, it is essential to investigate the roles and therapeutic implications of these dying signals in regulating cancer immune escape in the TME.

The TME plays a pivotal role in determining cancer chemoresistance and metastasis ^17^. Accumulating studies have demonstrated that chemotherapy can alter the TME and enhance immune responses in the TME ^18^. TAMs are the major tumor-infiltrating immune cell population within multiple solid tumors including breast cancer ^19^. Clinical evidence has revealed that TAM elevation usually predicates poor overall survival (OS) and clinical outcome in patients with breast cancer ^20^. TAMs are commonly regarded as the partners in the crime of tumor cells, promoting tumor immune escape, angiogenesis, growth, and metastasis ^19^. Notably, multiple chemotherapeutics, including paclitaxel, doxorubicin, and cyclophosphamide treatment, have been reported to stimulate breast cancer metastasis by generating a favorable pro-metastatic TME *via* increasing macrophage infiltration ^7, 21^. However, the molecular mechanisms underlying chemotherapy-induced TAM elevation are still largely unknown. As the clearance of dying cancer cells and dying signals are mainly attributed to macrophage phagocytosis, it is important to investigate how macrophages respond to chemotherapy-induced dying signals to remodel the pro-metastatic TME.

Exosomes are a subclass of membrane-coated vesicles that are 30–120 nm in diameter and are extracellularly secreted by exocytosis from cells. The exosomal content is heterogeneous, comprising proteins, DNAs, RNAs (mRNA, microRNA, and noncoding RNA), lipids, and metabolites. Exosomes have emerged as an ideal tool for early diagnosis, prognostic prediction, and therapeutic drug delivery of various cancers owing to their natural ability to mediate intercellular communication, as well as their high stability, endogenous origin, and low immunogenicity. Exosomes play a central role in the TME by mediating intercellular communication between cancer and stromal cells ^22^. Cancer cells use exosomes as a novel mechanism to transfer the malignant phenotype to normal fibroblasts and endothelial cells to establish a pro-metastatic TME. In addition, tumor cell-derived exosomes can inhibit the anti-cancer immune response by inducing the apoptosis of cytotoxic T cells, blocking the differentiation of monocytes into dendritic cells, and inducing the activation of immunosuppressive cells such as bone marrow suppressor cells (myeloid-derived suppressor cell, MDSC) and regulatory T cells (Tregs), which finally induce the immune escape of tumor cells ^23–25^. Notably, almost all of the existing exosome-related studies have focused on exosomes released from living tumor cells ^26, 27^. Cancer chemotherapy will inevitably lead to massive cancer cell death and therefore increase the release of exosomes from dying cancer cells. To the best of our knowledge, the existing knowledge of the biological effect of dying cancer cell-released exosomes on TME remodelling and tumor metastasis is limited. Keklikoglou *et al*. reported that neoadjuvant chemotherapy of breast cancer using taxanes and anthracyclines could elicit tumor-derived extracellular vesicles (EVs) with the enhanced pro-metastatic capacity. These EVs could induce Ly6C^+^CCR2^+^ monocyte expansion in the pulmonary pre-metastatic niche to facilitate the establishment of lung metastasis ^8^. Given the abundant infiltration and phagocytic nature of TAMs in the TME, it is interesting to investigate whether chemotherapy-induced dying cancer cell-released exosomes could affect the immune escape and metastasis of breast cancer by modulating TAMs in the TME.

In this study, we systematically demonstrated that exosomal CXCL1 signal, released from dying breast cancer cells following chemotherapy, could favor breast cancer immune escape and metastasis by transcriptionally activating TAM/PD-L1 signaling. In addition, TPCA-1 was screened as a promising small molecule to chemosensitize breast cancer by inhibiting dying cell-released exosomal CXCL1 signal.

## 2. Methods

### 2.1 Cell culture and induction

Breast cancer 4T1 cells (KG338) and Raw264.7 macrophages (KG240) were obtained from Nanjing KeyGen Biotech (Nanjing, China). 4T1-Luc cells were generated by transfecting 4T1 cells with the lentiviral luciferase reporter plasmid. For M1 and M2 macrophage induction, cells were stimulated with 100 ng/ml LPS, or 10 ng/ml IL-4 and 10 ng/ml IL-13 for 24 h, respectively. The identities of all these cell lines have been authenticated by short tandem repeat profiling.

### 2.2 Exosome isolation, quantification, observation, and particle size detection

For exo-dead isolation, 4T1 cells were cultured in the exosome-depleted medium and treated with 1 μM paclitaxel for 24 h to induce apoptosis. Then, cells were harvested and stained with Annexin V-FITC solution (70-APCC101-100, MultiSciences, Hangzhou, China) for 5 min. The Annexin V-FITC-positive cells were sorted by fluorescence-activated cell sorting technology using a FACS Aria III flow cytometer (BD Biosciences, Franklin Lakes, NJ, USA) and then cultured in the exosome-depleted medium for 24 h. Then, the cell culture medium was harvested to isolate exo-dead using Ribo^TM^ Exosome Isolation Reagent (C10130-2, Ribo Biotech, Guangzhou, China) or using a differential ultracentrifugation method ^28^. Additionally, exo-alive was isolated from the cell culture medium of untreated 4T1 cells. The protein concentration of exosomes was quantified by the BCA method and the structure was observed by a transmission electron microscope ^28^. The exosome particle sizes were detected using a Flow Nano Analyzer (NanoFCM Inc., Xiamen, China) ^29^.

### 2.3 Western blotting

Western blotting assay was conducted as previously reported ^30^. The following antibodies were used: Alix (12422-1-AP, RRID:AB_2162467, Proteintech, Wuhan, China), TSG101 (67381-1-Ig, RRID:AB_2882628, Proteintech), CD81 (66866-1-Ig, RRID:AB_2882203, Proteintech), Calnexin (ab133615, RRID:AB_2864299, Abcam, Cambridge, MA, USA), PD-L1 (DF6526, RRID:AB_2838488, Affinity, Changzhou, China), CXCL1 (AF5403, RRID:AB_2837887, Affinity), EED (85322T, RRID:AB_2923355, CST, MA, USA), and β-actin (4970S, RRID:AB_2223172, CST).

### 2.4 Zebrafish breast cancer xenotransplantation model

The zebrafish breast cancer xenotransplantation model was established as previously described ^31^. Briefly, 200 Dil-stained 4T1 cells were injected into the perivitelline space of each AB strain zebrafish embryo at 48 h post-fertilization using a microinjector. For the macrophage co-injection groups, 200 Dil-stained 4T1 cells and 600 Raw264.7 cells were co-injected. For exosome treatment, exosomes were co-injected with cells at the indicated doses. After treatment for 48 h, breast cancer growth and metastasis in zebrafish were observed under a Nikon SMZ25 stereomicroscope.

### 2.5 Animal experiments

The animal study was approved by the Institutional Animal Care and Use Committee of Guangdong Provincial Hospital of Chinese Medicine (No. 2021073). For 4T1-Luc xenograft establishment, 2×10^6^ 4T1-Luc cells were inoculated subcutaneously into the mammary fat pads of female BALB/c mice (6 weeks old). The animals were randomly grouped using a random number table. To investigate the effect of exo-dead on breast cancer growth and metastasis, 4T1-Luc xenograft-bearing mice were randomly divided into the saline group (peritumoral injection with saline solution, 200 μl/20 g weight, q3d) and exo-dead group (peritumoral injection with exo-dead, 200 μg/20 g weight, q3d). To investigate the effect of exosomal CXCL1 overexpression on breast cancer growth and metastasis, 4T1-Luc xenograft-bearing mice were randomly divided into three groups, including saline group, exo-dead group, and exo-dead^rCXC1^ group (peritumoral injection with exo-dead^rCXC1^, 200 μg/20 g weight, q3d). To investigate the effects of exosomal CXCL1 knockdown or macrophage deletion on the pro-tumor activity of exo-dead, 4T1-Luc xenograft-bearing mice were randomized into four groups, including saline group, exo-dead group, exo-dead^shCXCL1^ group (peritumoral injection with exo-dead^shCXC1^, 200 μg/20 g weight, q3d), and exo-dead + CL group (combined treatment with exo-dead and clodronate liposomes, CL). Subsequently, 200 μl CL (40337ES10, Yeasen Biotech, Shanghai, China) was injected intraperitoneally into 4T1-Luc xenograft-bearing mice 2 days before the exosome injection, following which, intraperitoneal injection (100 μl per mouse) was continued once every 2 weeks during the animal experiment period. To investigate the combined effect of TPCA-1 and paclitaxel treatment, 4T1-Luc xenograft-bearing mice were randomly divided into four groups, including saline group, TPCA-1 group (10 mg/kg/d, intraperitoneal injection), paclitaxel group (10 mg/kg/3d, intraperitoneal injection), and TPCA-1 + paclitaxel group. To investigate the effect of TPCA-1 treatment on the pro-tumor activity of exo-dead, 4T1-Luc xenograft-bearing mice were randomly divided into four groups: saline group, exo-dead group, exo-dead + TPCA-1 group, and exo-dead^rCXCL1^ + TPCA-L1 group. The administration doses of exo-dead, exo-dead^rCXCL1^, or TPCA-L1 were the same as those stated above. The mice were imaged using an IVIS Lumina XR *in vivo* imaging system (PerkinElmer, MA, USA) to monitor tumor growth and metastasis. At the end of the animal experiment, mice were euthanized and the primary cells were isolated from tumors and subjected to macrophage phenotypic analysis as indicated below. The blood samples were subjected to biochemical analysis as previously reported ^32^ to detect the hepatotoxicity, nephrotoxicity, or hematotoxicity of TPCA-1, and HE staining assay was applied as previously reported ^31^ to investigate the lung metastasis differences in mice in different groups.

### 2.6 Macrophage phenotype, population, and PD-L1 expression analyses

For the phenotype analysis of Raw264.7 macrophages, cells were first treated as indicated. Then, macrophages were harvested and incubated with FITC-conjugated F4/80 antibody (SC-71085, Santa Cruz, CA, USA), PE-conjugated CD206 antibody (141705, Biolegend, CA, USA), or PE-Cy7-conjugated CD206 antibody (E-AB-F1135H, Elabscience, Houston, TX, USA) for 30 min at 37°C. For the phenotypic analyses of primary macrophages isolated from mouse 4T1-Luc xenografts, cells were incubated with CD45-PE-Cy7 (25-0451-82, eBioscience, Waltham, MA, USA), F4/80-APC antibody (17-4801-82, eBioscience), and CD206-PE antibody (141705, Biolegend) for 30 min at 37°C. For PD-L1 expression analyses of primary macrophages, cells were incubated with CD45-PE-Cy7 (25-0451-82, eBioscience), F4/80-FITC antibody (SC-71085, Santa Cruz), and PD-L1-APC antibody (124312, Biolegend) for 30 min at 37°C. After incubation, the cells were washed once with PBS and subjected to flow cytometry analysis.

### 2.7 Macrophage phagocytosis assay

Briefly, exosomes were labelled with the PKH67 green fluorescent cell linker (2 μM, MINI67, Sigma-Aldrich, Missouri, USA). The labelling process was stopped by adding serum to the mixture. Then, Raw264.7 cells were treated with PKH67-labeled exosomes for 1–4 h. After washing with PBS three times, Raw264.7 cells were harvested and subjected to flow cytometry to measure the green fluorescence intensities of the cells. To visualize the phagocytosis process of exosomes by macrophages, the PKH67-treated macrophages were fixed, permeabilized, and then incubated with ActinRed (5 U/ml, KGMP0012, KeyGEN) for 20 min. Lastly, the cells were observed using an LSM710 confocal microscope (Zeiss, Oberkochen, Germany).

### 2.8 Co-culture of breast cancer cells and macrophages

The six- or 24-well Transwell co-culture system was used for the co-culture of breast cancer cells and macrophages. In brief, Transwell inserts were placed in six- or 24-well culture plates. Breast cancer cells and macrophages were seeded into different Transwell chambers. The Transwell inserts were separated by a 0.4-μm permeable membrane that allowed the free exchange of media and soluble molecules. For the Transwell assay, an 8-μm pore size Transwell chamber was used to allow the migration of breast cancer cells.

### 2.9 Cell counting, colony formation, wound healing, Transwell, and CCK-8 assays

Cell counting and colony formation assays were conducted ^30^ to investigate the proliferation and colony formation abilities of breast cancer cells, respectively. Wound healing and Transwell assays were conducted ^31^ to investigate the migration and invasion abilities of breast cancer cells in the co-culture system when they were treated as indicated. CCK-8 assay was conducted ^31^ to investigate the viability of breast cancer cells in the co-culture system when they were treated with exo-alive, exo-dead, TPCA-1, paclitaxel, or their combination.

### 2.10 Stem cell population analysis and mammosphere formation assay

Stem cell population analysis was conducted as previously described ^30^ using the ALDEFLUOR Stem Cell Identification Kit (No.01700, STEMCELL, Cambridge, MA, USA) and NovoCyte flow cytometer (ACEA, Hangzhou, China), and analyzed using NovoExpress. Mammosphere formation assay was conducted as previously described ^30^ to investigate the stemness changes of breast cancer cells following treatment. The number of mammospheres was quantified microscopically.

### 2.11 Mouse chemokine array assay

The Mouse Chemokine Array C1 (AAM-CHE-1-4, RayBio, Norcross, GA, USA) was used to analyze the chemokine composition differences between exo-alive and exo-dead. Briefly, the chemokine antibody-coated membranes were blocked with blocking buffer for 30 min and incubated with equal amounts of exo-alive or exo-dead solution overnight at 4 °C. Then, the membranes were washed and incubated with the biotin-conjugated detection antibody cocktail for 2 h, and the HRP-conjugated streptavidin for 1 h. Finally, the membranes were washed, subjected to chemiluminescence, developed, and photographed.

### 2.12 CXCL1 secretion inhibitor screening

To screen the potential CXCL1 secretion inhibitor of 4T1 cells from the Chemokine Inhibitor Library (L7600, TOPSCIENCE, Shanghai, China), 4T1 cells were treated with 80 types of chemokine inhibitors (1 μM) for 48 h. Subsequently, the concentration of CXCL1 in cell culture supernatants was detected using the Mouse CXCL1 ELISA Kit.

### 2.13 ELISA

An ELISA was conducted to detect the CXCL1 concentrations in different exosome preparations. Briefly, equal quantities of exosomes were lysed by RIPA and sonication. CXCL1 concentrations in the lysed exosomes were detected using the Mouse CXCL1 ELISA Kit (SEA041Mu, USCN Business, Wuhan, China) as previously described ^31^.

### 2.14 Immunofluorescence assay

Immunofluorescence analysis was conducted as previously described ^30^. The following primary antibodies were used in the immunofluorescence assay including PD-L1 (DF6526, Affinity), CXCL1 (AF5403, Affinity), Flag (M185-3L, MBL International Corporation, Woburn, MA, USA), CD206 (141704, Biolegend), PD-L1 (66248-1-IG, Proteintech), and EED (DF7308, Affinity) antibodies. The following secondary antibodies were used in the immunofluorescence assay: Alexa Fluor® 555 conjugated-anti-rabbit IgG (no.4413S, CST), Alexa Fluor® 488 conjugated-anti-rabbit IgG (4412S, CST), Alexa Fluor® 555 conjugated-anti-mouse IgG (A21422, ThermoFisher, Waltham, MA, USA), FITC conjugated-anti-Rat IgG (SC-2011, Santa Cruz), and Alexa Fluor® 488 conjugated-anti-mouse IgG (4408S, CST). Fluorescence images were obtained using an LSM710 confocal microscope.

### 2.15 Transfection of plasmid and siRNA

A commercialized CXCL1 recombinant plasmid with a C-terminal FLAG tag was purchased from Dahong Biosciences (Guangzhou, China). The shRNA plasmids for CXCL1, EED, and PD-L1 as well as the PD-L1 recombinant plasmid were purchased from Vigene Biosciences (Jinan, China). Plasmids were transfected into the indicated cells using the LipoFiter^TM^ reagent (Hanbio Biotech, Shanghai, China) ^30^.

### 2.16 Circulating tumor cell (CTC) detection

The number of CTCs in the blood of tumor-bearing mice was measured by detecting the relative expression levels of the luciferase gene derived from breast cancer 4T1-Luc cells. Genomic DNA was extracted from mouse peripheral blood and measured by QPCR assay using the following primers: 5′-GCTCAGCAAGGAGGTAGGTG-3′ (forward) and 5′-TCTTACCGGTGTCCAAGTCC-3′ (reverse) for luciferase; and 5′-GGAGGGGGTTGAGGTGTT-3′ (forward) and 5′-GTGTGCACTTTTATTGGTCTCAA-3′ (reverse) for mouse *β-actin*.

### 2.17 QPCR

QPCR was conducted as previously described ^30^. The primer sequences were as follows: 5′-GCTCCAAAGGACTTGTACGTG-3′ (forward) and 5′-TGATCTGAAGGGCAGCATTTC-3′ (reverse) for mouse *PD-L1*.

### 2.18 Double luciferase reporter gene assay

The double luciferase reporter gene assay was conducted using the Secrete-Pair™ Dual Luminescence Assay Kit (LF031, Genecopeia, Rockville, MD, USA) ^31^ to investigate the *PD-L1* promoter activity changes of Raw264.7 cells when treated as indicated. The *PD-L1* promoter plasmid (MPRM25392-PG04, Genecopeia) was transfected into Raw264.7 cells using Vigenefection (FH880806, Vigene Biosciences)^31^.

### 2.19 DNA-pull down-MS

The DNA-pull down-MS assay was conducted by Huijun Biotechnology (Guangzhou, China) to identify the transcription factor responsible for the CXCL1-induced promoter activity of *PD-L1*. Briefly, the biotinylated promoter fragment of *PD-L1* was synthesized by PCR using the following primers: 5′-TCTTGAACGGCAAGACAAC-3′ (forward) and bio-5′-TTCTGACCCAGCTACCTAC-3′ (reverse). The pull-down experiments were conducted as previously described ^33^ using the biotinylated *PD-L1* promoter fragment. The purified proteins underwent MS analysis by Huijun Biotechnology. The enriched protein was obtained by comparing the identified proteins with the control group.

### 2.20 Chromatin immunoprecipitation (CHIP)-PCR

CHIP assay was conducted by immune-precipitating the DNA fragments with EED antibody (85322T, CST) using the CHIP Assay Kit (P2078, Beyotime, Shanghai, China) ^34^. Analysis of the genomic sequence of the *PD-L1* promoter (NC_000085.7:29342838-29344837) revealed a potential binding site (5’-GTTCCACTC-3’, site: –437 to –429 bp) for the transcription factor EED. This region in the immune-precipitated DNA samples was amplified by PCR assay using the following primers: 5′-AAGGTGGGAGCTGTAGAGGAA-3′ (forward) and 5′- TGCTACTGAGAGGCTGTCGAT-3′ (reverse).

### 2.21 Statistical analysis

Student’s t-test and one-way ANOVA were used for comparisons among groups. Levene’s Test of Equality of Variances was used to assess the assumption of homogeneity of variance. Survival curves were calculated using Kaplan–Meier analysis and were compared using the log-rank test. Data are represented as the mean ± SD. All tests were 2-sided and *P* < 0.05 was considered statistically significant.

## 3. Results

### 3.1 Exosomes from dying breast cancer cells (exo-dead) induce breast cancer lung metastasis and promote TAM infiltration *in vivo*

Paclitaxel remains one of the most commonly used chemotherapeutics for breast cancer treatment ^3^. To isolate exo-dead, breast cancer 4T1 cells were cultured in exosome-depleted medium, and paclitaxel was used to induce apoptosis of 4T1 cells. Subsequently, the early apoptotic populations were sorted by flow cytometry. Exo-dead and exo-alive were isolated from the supernatants of apoptotic 4T1 cells and untreated 4T1 cells, respectively. For exosome characterization, it was found that the isolated exosomes exhibited a lipid bilayer structure and had a median diameter of 79.0 nm for exo-dead and 83.3 nm for exo-alive. Both exosomes also exhibited elevated expression of exosome positive markers including Alix, TSG101, and CD81, while exhibiting little expression of the exosome negative marker calnexin (**Figure 1A**). These results suggest the successful isolation of both exo-alive and exo-dead. Additionally, quantitative analysis results showed that paclitaxel treatment significantly elevated the number of exosomes secreted from 4T1 cells (**Figure 1B**). Next, the effects of exo-dead and exo-alive on breast cancer growth and metastasis were investigated *in vitro* and *in vivo*. Both exo-alive and exo-dead treatment had little effect on the proliferation of breast cancer cells *in vitro* and in the zebrafish breastcancer xenotransplantation model *in vivo* (**Figure 1-figure supplement 1**). Macrophages are considered as the most abundant immune cell subset in the TME of breast cancer ^19^. Additionally, macrophages have a powerful ability to phagocytize foreign bodies, while phagocytosis also represents an efficient way for exosome uptake. Interestingly, further investigations found that exo-dead (50–100 μg/ml) treatment significantly promoted the metastasis of 4T1 cells in the presence of Raw264.7 macrophage co-injection, which was not significantly achieved by exo-alive (**Figure 1C**). More importantly, peritumoral injection with exo-dead significantly promoted the growth and lung metastasis of mouse breast cancer 4T1-Luc xenografts (**Figure 1D–E**), which was accompanied by the elevated infiltration of CD45^+^/F4/80^+^/CD206^+^ TAMs in the TME (**Figure 1F**). Meanwhile, exo-dead treatment had no significant effect on the body weights of mice (**Figure 1D**). These results suggest that exo-dead may promote breast cancer growth and metastasis by modulating macrophage polarization in the TME. Taken together, these findings show that dying breast cancer cells-released exosomes induce breast cancer growth and lung metastasis, as well as elevating TAM infiltration *in vivo*.

**Figure 1.**
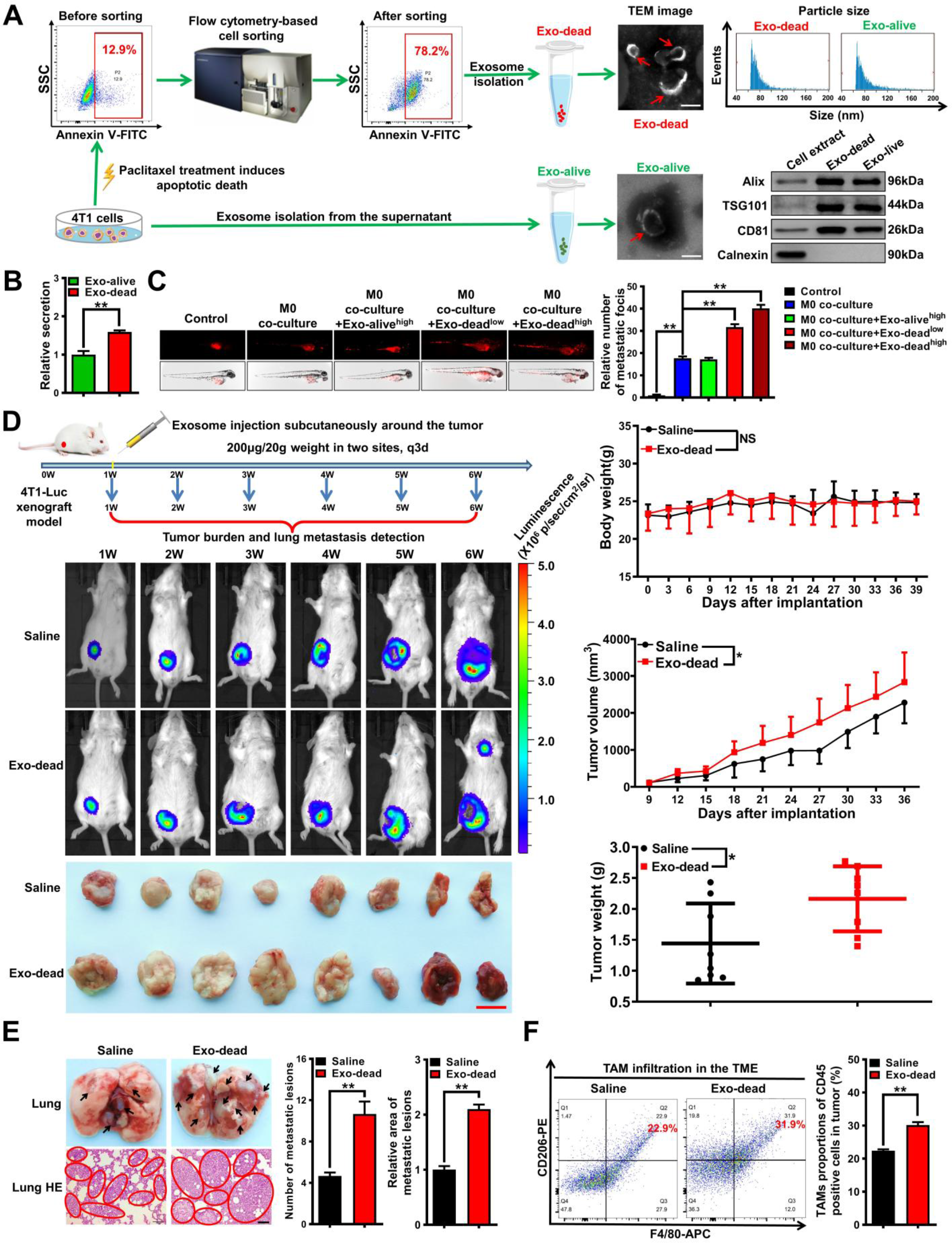
Exo-dead induces breast cancer lung metastasis and promotes TAM infiltration *in vivo*. **(A)** Diagram of the exo-dead and exo-alive separation procedures and their representative transmission electron microscopy (TEM) images. Scale bar: 100 nm. The sizes of the exosome particles were detected using a Flow Nano Analyzer, and their protein markers were identified by western blot analysis. **(B)** The BCA method was conducted to detect the relative secretion content of exo-dead and exo-alive; n = 3. **(C)** Representative images of zebrafish breast cancer xenotransplantation model assay. The effects of exo-alive (100 μg/ml) and exo-dead (50–100 μg/ml) on the metastasis of 4T1 cells in the presence or absence of M0 co-injection were investigated; n= 6. **(D)** Schematic diagram of the animal assay and representative pictures of the *in vivo* imaging assay and tumors. Peritumoral injection with exo-dead (200 μg/20 g weight, q3d) promoted the growth of 4T1-Luc xenografts in terms of both tumor volume and weight, but had no significant effect on the weight of the mice; n = 8. Scale bar: 1 cm. **(E)** Representative images of the lungs and the lung HE staining assay. Metastatic foci were identified by HE staining of the lung sections. Scale bar: 100 μm; n= 3. **(F)** Infiltration levels of CD45^+^/F4/80^+^/CD206^+^ TAMs in mammary tumors; n= 3. \**p* < 0.05, \*\**p* < 0.01.

### 3.2 Exo-dead promotes the metastasis and chemoresistance of breast cancer cells by inducing macrophage M2 polarization

Next, we investigated whether exo-dead induces breast cancer growth and metastasis by modulating TAMs *in vitro*. First, immunofluorescence and flow cytometry analysis showed that PKH67-labeled exo-dead could be time-dependently phagocytosed by Raw264.7 macrophages (**Figure 2A**). Undifferentiated macrophages (M0) can polarize into pro-inflammatory M1 phenotype or anti-inflammatory M2 phenotype under different stimulations. Flow cytometry results showed that exo-dead treatment significantly promoted M0 macrophage polarization into the M2 phenotype in a dose-dependent manner *in vitro* (**Figure 2B**). Accumulating studies have suggested that TAMs can promote cancer growth and metastasis. Therefore, breast cancer 4T1 cells and Raw264.7 macrophages were co-cultured *in vitro* using the Transwell model to simulate their coexistence (**Figure 2C**). It was found that exo-dead administration (25–100 μg/ml) in the culture medium of Raw264.7 macrophages could significantly promote the proliferation (**Figure 2C**), colony formation (**Figure 2D**), migration, and invasion abilities (**Figure 2E**) of the co-cultured breast cancer 4T1 cells in a concentration-dependent manner. Meanwhile, exo-dead treatment reduced the chemosensitivity of 4T1 cells to paclitaxel (**Figure 2F**). Breast cancer stem cells (BCSCs) are considered as the root of breast cancer metastasis and chemoresistance. Exo-dead treatment also elevated the subpopulation of ALDH^+^ BCSCs in 4T1 cells (**Figure 2G**), and enhanced their mammosphere formation abilities (**Figure 2H**). Taken together, these results indicate that exo-dead promotes the metastasis and chemoresistance of breast cancer cells by inducing macrophage M2 polarization.

**Figure 2.**
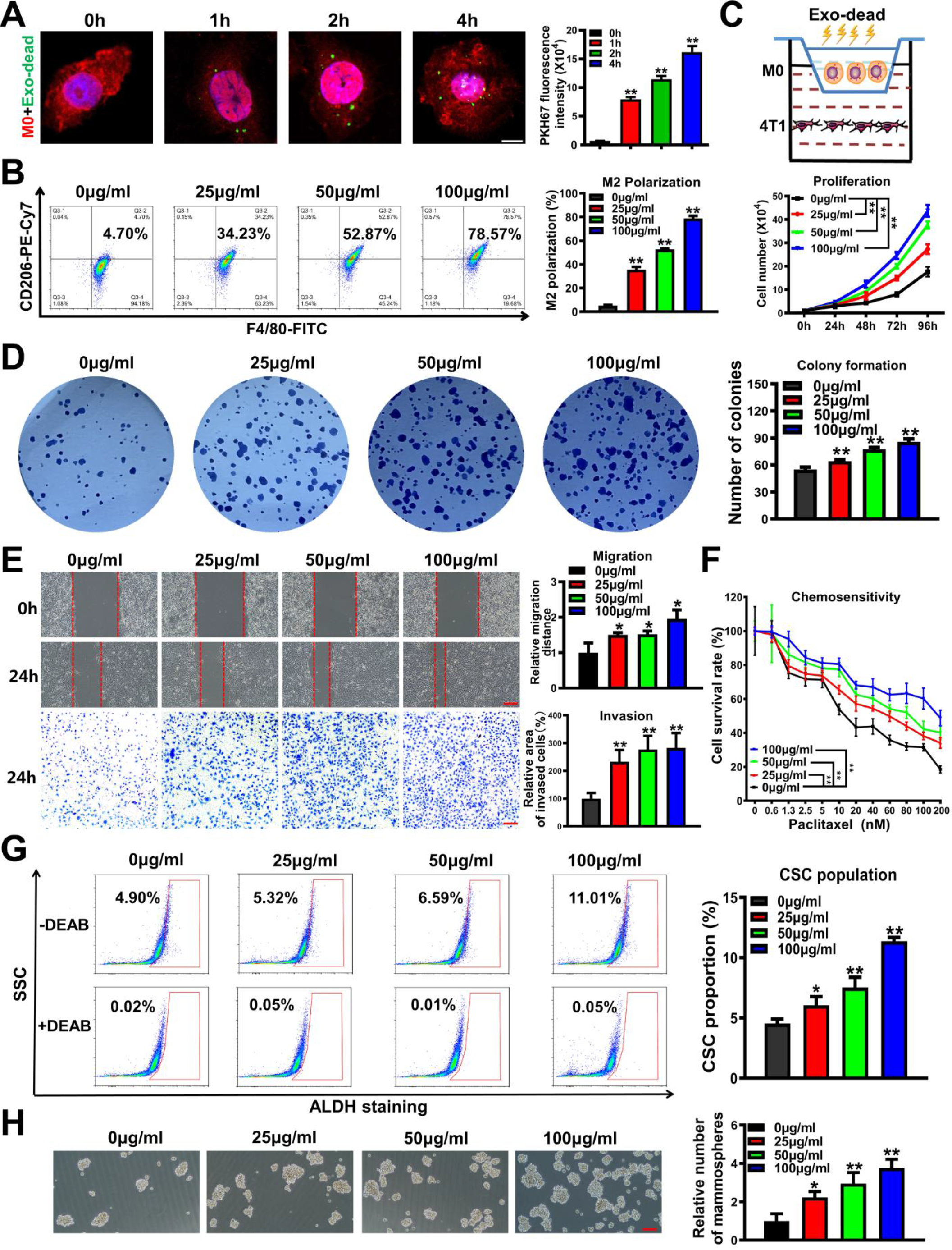
Exo-dead promotes the metastasis and chemoresistance of breast cancer cells by inducing macrophage M2 polarization. **(A)** Exo-dead uptake by Raw264.7 macrophages was visualized using immunofluorescence labeling and quantified by flow cytometry. Exo-dead was labeled with PKH67 (green). Raw264.7 macrophages were labeled with ActinRed (red) and DAPI (blue); n = 3. Scale bar: 5 μm. **(B)** The polarization changes of Raw264.7 macrophages after exo-dead treatment (25–100 μg/ml) for 48 h; n= 3. **(C–D)** Diagram of 4T1 and Raw264.7 cell co-culture using the Transwell system. Changes in proliferation (C) and colony formation ability (D) of the co-cultured 4T1 cells after exo-dead treatment; n = 3. **(E)** Migration and invasion efficacy changes of the co-cultured 4T1 cells after exo-dead treatment. Scale bars: 200 μm; n = 3. **(F)** Changes in chemotherapeutic sensitivity of the co-cultured 4T1 cells to paclitaxel after exo-dead treatment for 48 h, as determined by CCK-8 assay; n = 8. **(G–H)** The ALDH^+^ BCSC subpopulation (G) and their mammosphere formation abilities (H) after exo-dead treatment for 48 h. Diethylaminobenzaldehyde (DEAB) is a specific inhibitor of ALDH activity. Scale bar: 200 μm; n = 3. \**p* < 0.05, \*\**p* < 0.01.

### 3.3 CXCL1^exo-dead^ induces macrophage M2 polarization by activating PD-L1 expression

Next, we sought to determine the bioactive molecule in exo-dead that is responsible for inducing macrophage M2 polarization. As emerging evidence has suggested that chemokines play important roles in inducing the activation and polarization of macrophages ^35^, we aimed to characterize the abundant chemokines contained in exo-dead using a chemokine array. As shown in **Figure 3A**, exo-dead contained multiple chemokines, among which, CXCL1 was the most abundant. More importantly, both the semiquantitative analysis of the chemokine array and quantitative ELISA confirmed that CXCL1 was significantly more upregulated in exo-dead compared to that in exo-alive. It has been reported that CXCL1 expression is significantly correlated with metastasis and poor OS in patients with breast cancer^36^. Therefore, we next investigated whether CXCL1 in exo-dead was responsible for the pro-metastatic effect of exo-dead. As shown in **Figure 3B**, CXCL1 knockdown in exo-dead or CXCL1 neutralizing antibody (NA) partially abrogated the induction effect of exo-dead on the M2 polarization of macrophages, leading to the decreased invasion of breast cancer cells in the co-culture system (**Figure 3C**). Increasing studies have suggested that macrophages represent the major cellular source for maintaining PD-L1 expression in the TME, while PD-L1 is crucial for the activation and M2 polarization of macrophages ^37, 38^. Here, western blotting revealed that exo-dead could induce PD-L1 expression in macrophages in a time-and dose-dependent manner (**Figure 3D**), while CXCL1 knockdown in exo-dead or CXCL1 NA administration inhibited PD-L1 expression (**Figure 3E–F**). Meanwhile, it was found that exo-dead induced macrophage M2 polarization by activating PD-L1 expression, as shown by the finding that PD-L1 knockdown partially abrogated the M2 polarization of macrophages induced by exo-dead (**Figure 3G**). Taken together, these findings show that CXCL1 is a crucial chemokine in exo-dead, which functions to mediate macrophage M2 polarization by activating PD-L1 expression.

**Figure 3.**
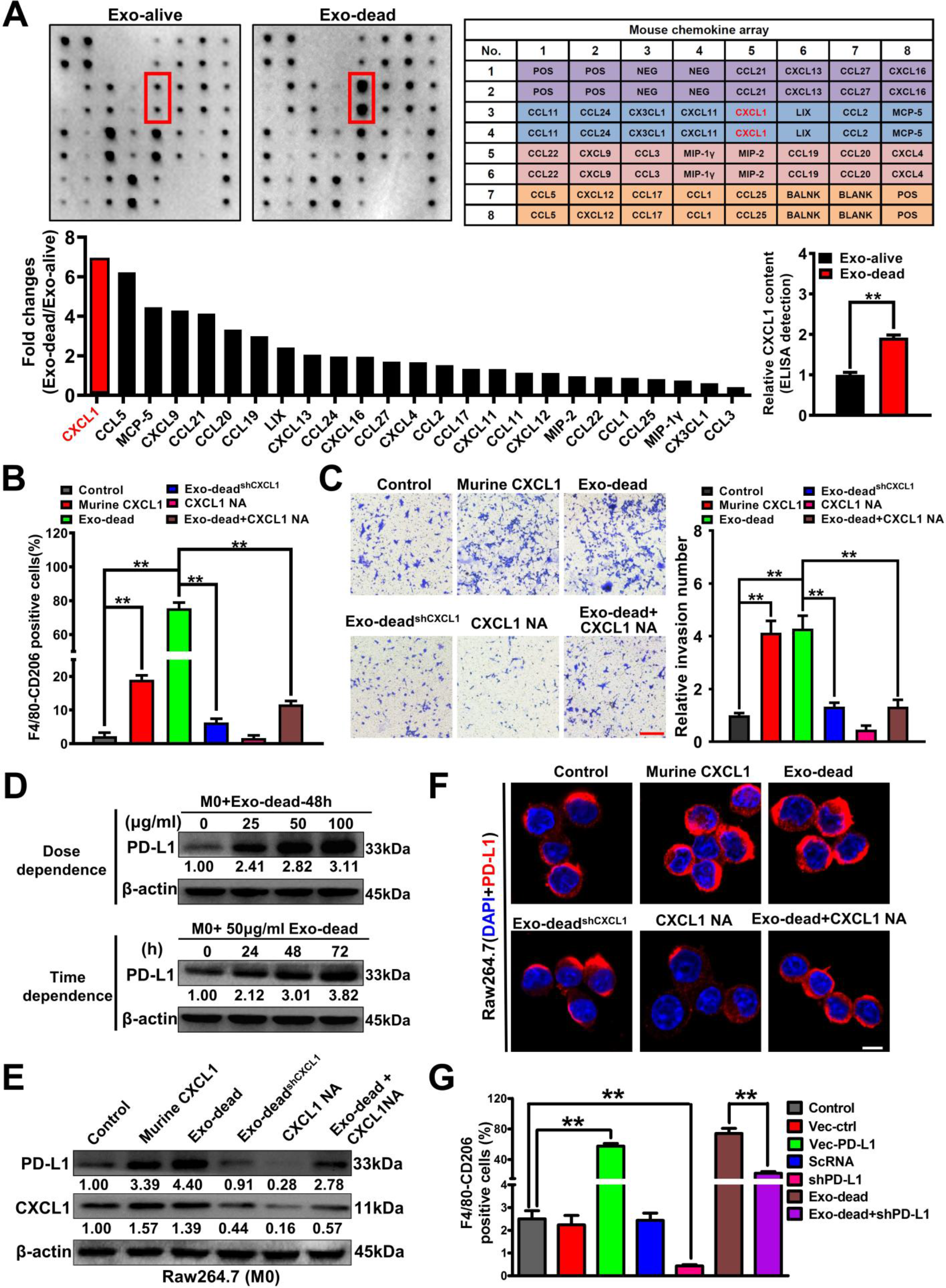
CXCL1^exo-dead^ induces macrophage M2 polarization by activating PD-L1 expression. **(A)** Chemokine array assay was conducted to characterize the differences in chemokine content between exo-dead and exo-alive. An ELISA was conducted to compare the relative CXCL1 content in exo-dead and exo-alive. **(B)** Changes in M2 phenotype polarization of Raw264.7 macrophages when treated with 10 ng/ml murine CXCL1, 50 μg/ml exo-dead, 50 μg/ml exo-dead^shCXCL1^, 5 μg/ml CXCL1 neutralizing antibody (NA), or exo-dead and CXCL1-NA combination for 48 h. **(C)** Representative images of Transwell assay. Raw264.7 macrophages were treated as indicated for 48 h and then co-cultured with 4T1 cells. Scale bar: 200 μm. **(D–F)** Expression changes of CXCL1 and PD-L1 in Raw264.7 macrophages when treated as indicated for 48 h. Scale bar: 10 μm. **(G)** The results of flow cytometry assay suggested that 50 μg/ml exo-dead treatment for 48 h induced the M2 polarization of Raw264.7 macrophages by activating PD-L1 expression; n = 3. \*\**p* < 0.01.

### 3.4 CXCL1^exo-dead^ promotes breast cancer growth and lung metastasis *in vivo* by activating TAM/PD-L1 signaling

Next, we sought to validate whether CXCL1 in exo-dead is crucial in inducing breast cancer growth and lung metastasis *in vivo*. To achieve this, Flag-tagged CXCL1 was overexpressed in 4T1 cells and exo-dead^rCXCL1-Flag^ was isolated from the supernatants of apoptotic 4T1/rCXCL1^Flag^ cells following paclitaxel treatment. As shown in **Figure 4A**, immunoblotting and immunofluorescence assays validated the successful generation of 4T1/rCXCL1^Flag^ cells. Meanwhile, ELISA confirmed that CXCL1 was significantly more elevated in exo-dead^rCXCL1^ compared to exo-dead (**Figure 4B**). It was also found that peritumoral injection with exo-dead^rCXCL1^ significantly accelerated the growth and lung metastasis of breast cancer in the mouse 4T1-Luc xenograft models compared to that of exo-dead (**Figure 4C–D**), which also resulted in decreased OS of the mammary tumor-bearing mice. More importantly, the tumor tissue immunofluorescence experiment clearly showed that Flag-tagged CXCL1 was predominantly phagocytosed by CD206^+^ TAMs in the TME (**Figure 4E**), which promoted PD-L1 expression and subsequent M2 polarization (**Figure 4E–F**). CTCs are a rare population of tumor cells that contribute to the development of metastatic disease following their release into the peripheral circulation from primary tumor sites. In this study, peripheral CTCs were detected by QPCR analysis using primers directed to the luciferase genes of 4T1-Luc cells. It was found that exo-dead^rCXCL1^ injection increased CTCs by 4.1-fold while exo-dead injection elevated CTCs by 2.3-fold (**Figure 4G**), suggesting an increased metastatic potential of breast cancer cells induced by exo-dead^rCXCL1^ compared to exo-dead. Taken together, these findings indicate that CXCL1^exo-dead^ promotes the growth and lung metastasis of breast cancer by activating TAM/PD-L1 signaling.

**Figure 4.**
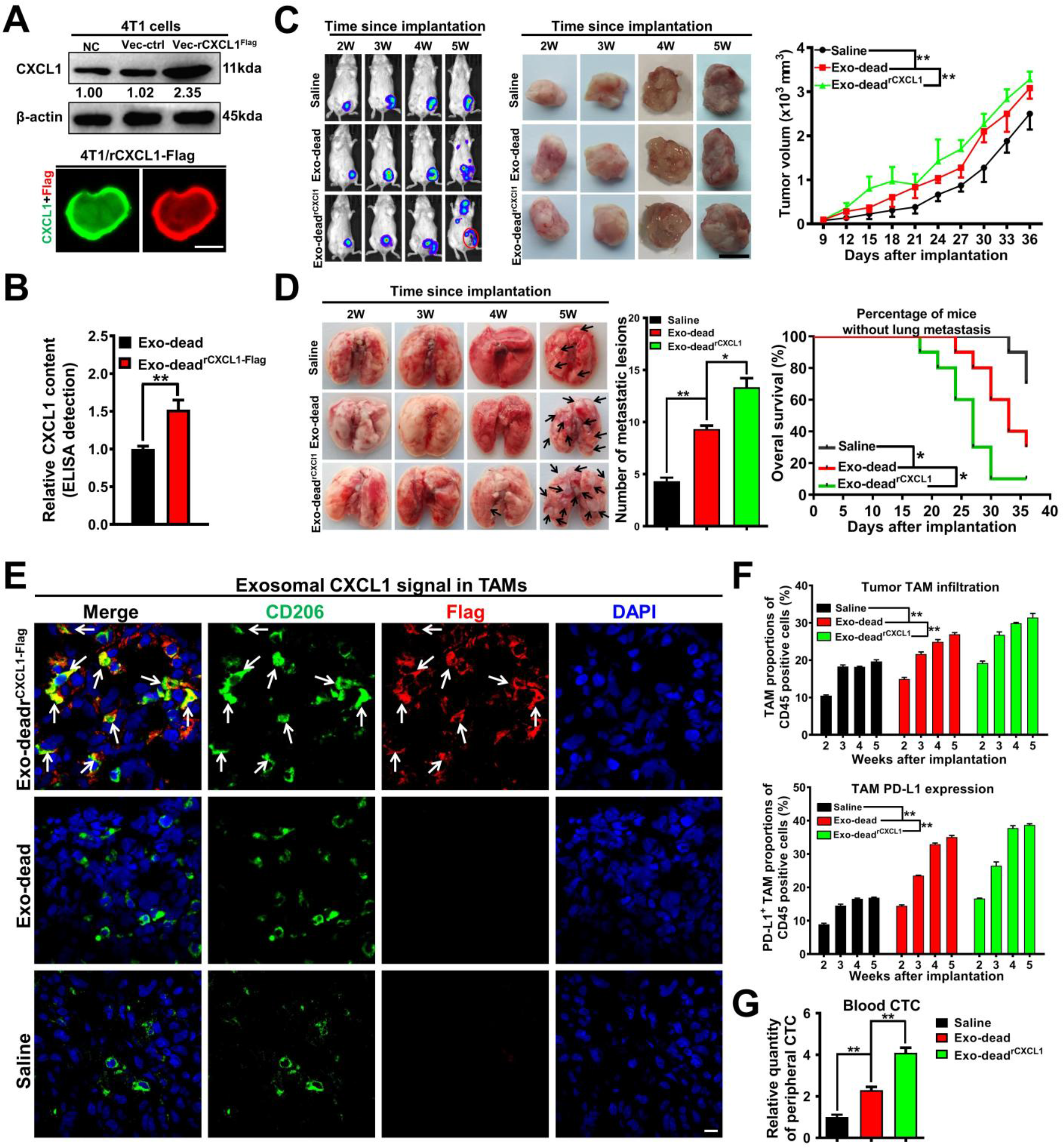
CXCL1^exo-dead^ promotes breast cancer growth and lung metastasis *in vivo* by activating TAM/PD-L1 signaling. **(A)** The successful generation of 4T1/rCXCL1^Flag^ cells was validated by immunoblotting and immunofluorescence assays. Scale bar: 5 μm. **(B)** The difference in CXCL1 content between exo-dead and exo-dead^rCXCL1-Flag^ was compared by ELISA. Exo-dead^rCXCL1-Flag^ was isolated from the supernatants of apoptotic 4T1/rCXCL1^Flag^ cells induced by paclitaxel treatment; n = 3. **(C–D)** Peritumoral injection with exo-dead^rCXCL1^ (200 μg/20 g weight, q3d) significantly accelerated breast cancer growth (C) and lung metastasis (D) compared to that of the exo-dead group (200 μg/20 g weight, q3d); Tumor volume: n = 6; Number of metastatic lesions, n=3; K-M curves of lung metastasis time, n=10. Scale bar: 1 cm. **(E)** Tumor tissue immunofluorescence experiment showed that flag-tagged CXCL1 (red) from exo-dead^rCXCL1-Flag^ was predominantly phagocytosed by CD206^+^ macrophages (green) in the TME. Scale bar: 5 μm. **(F)** The infiltration levels of CD45^+^/F4/80^+^/CD206^+^ TAMs (up) and CD45^+^/F4/80^+^/PD-L1^+^ TAMs (down) in the TME of mice following treatment with saline, exo-dead, or exo-dead^rCXCL1-Flag^; n = 3. **(G)** QPCR assay was conducted to investigate the CTC quantity in the peripheral blood of mice following treatment with saline, exo-dead, or exo-dead^rCXCL1-Flag^; n = 3. \**p* < 0.05, \*\**p* < 0.01.

### 3.5 Exosomal CXCL1 knockdown or macrophage depletion inhibits exo-dead-induced breast cancer growth and lung metastasis *in vivo*

Next, CXCL1 knockdown in exo-dead or macrophage depletion in the TME was applied to further validate the crucial roles of exosomal CXCL1 and TAMs in exo-dead-induced breast cancer growth and lung metastasis. To achieve this, exo-dead^shCXCL1^ was isolated from the supernatants of apoptotic 4T1/shCXCL1 cells induced by paclitaxel treatment. ELISA showed that the CXCL1 level in exo-dead^shCXCL1^ was significantly downregulated compared to that in exo-dead (**Figure 5A**). Macrophages in the TME were specifically depleted by CL treatment. As shown in **Figure 5B–C**, both CXCL1 knockdown in exo-dead and macrophage depletion partially inhibited the promotion effect of exo-dead on breast cancer growth and lung metastasis in the mouse 4T1-Luc xenograft model, leading to the increased OS of the mammary tumor-bearing mice. Meanwhile, both CXCL1 knockdown in exo-dead and macrophage depletion remarkably decreased the intratumoral infiltration levels of CD45^+^/F4/80^+^/CD206^+^ TAMs (**Figure 5D**) and CD45^+^/F4/80^+^/PD-L1^+^ TAMs induced by exo-dead (**Figure 5E–F**), which was also accompanied by decreased CTCs in the blood of 4T1-Luc xenograft-bearing mice (**Figure 5G**). These results indicated that exosomal CXCL1 played an important role in recruiting TAMs and activating their PD-L1 expression. Taken together, these findings demonstrate that CXCL1 is the crucial bioactive chemokine in exo-dead to induce breast cancer growth and metastasis, while macrophages act as the essential target cells of exo-dead in this biological process.

**Figure 5.**
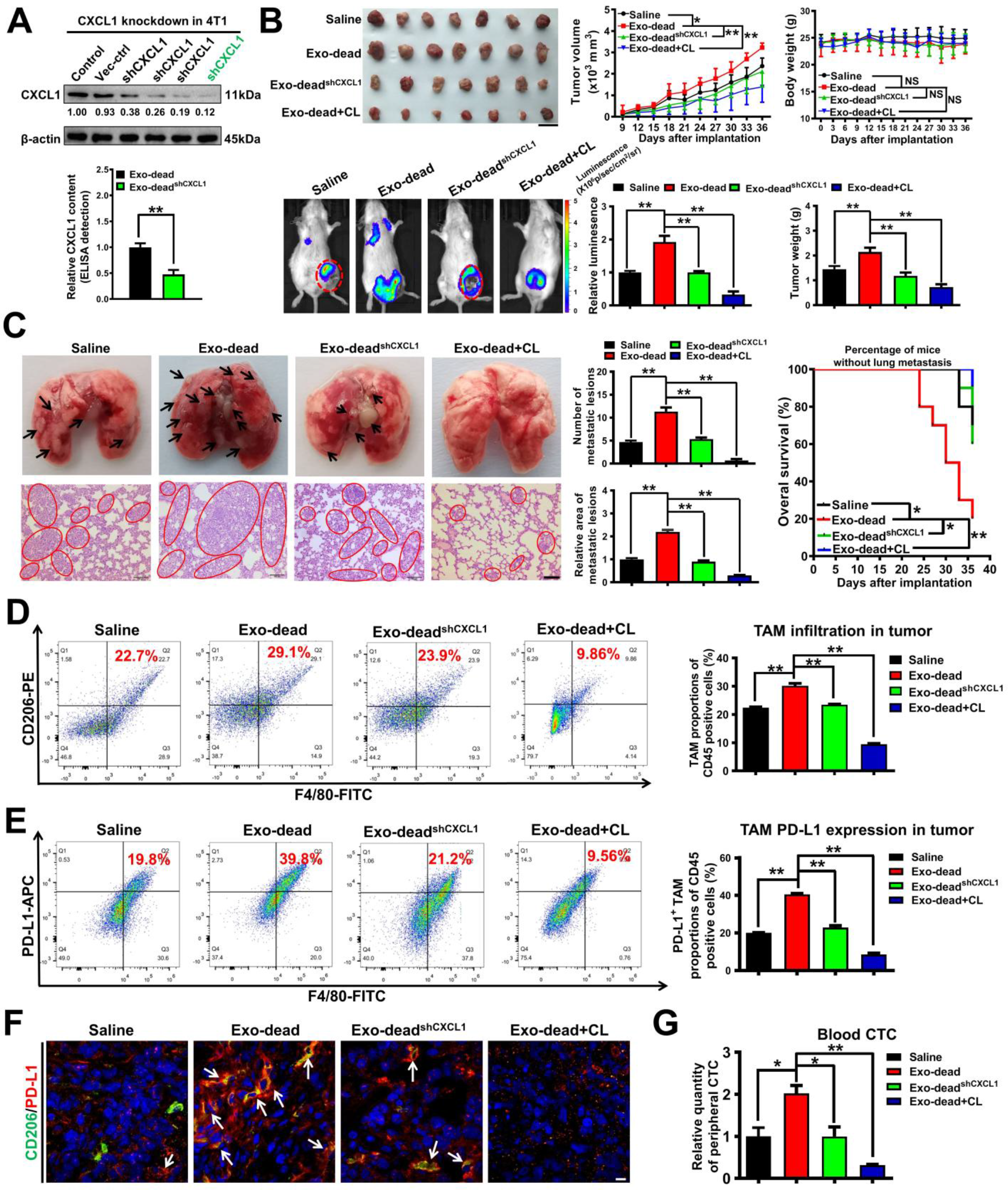
Exosomal CXCL1 knockdown or macrophage depletion inhibits exo-dead-induced breast cancer growth and lung metastasis *in vivo*. **(A)** The successful generation of 4T1/shCXCL1 cells was verified by western blotting assay. The difference in CXCL1 content between exo-dead and exo-dead^shCXCL1^ was compared by ELISA. Exo-dead^shCXCL1^ was isolated from the supernatants of apoptotic 4T1/shCXCL1 cells induced by paclitaxel treatment; n = 3. **(B)** Representative images of the tumors (n = 7) and *in vivo* imaging assay (n = 3), and mouse weight and tumor volume curves (n = 7). Clodronate liposomes (CL) were used to deplete macrophages in the TME of the 4T1-Luc xenograft model. Scale bar: 2 cm. **(C)** Representative images of the lungs (n = 3) and lung HE assay (n = 3) as well as the K-M curves of lung metastasis time (n = 10). Scale bar: 100 μm. **(D–E)** The infiltration levels of CD45^+^/F4/80^+^/CD206^+^ TAMs (D) and CD45^+^/F4/80^+^/PD-L1^+^ TAMs (E) in the TME of mice following treatment with exo-dead, exo-dead^shCXCL1^, or the combination of exo-dead and CL; n = 3. **(F)** CD206 (green) and PD-L1 (red) expression levels in the TME. Arrows indicate PD-L1 expression in TAMs. Scale bar: 10 μm. **(G)** The quantity of CTCs in the peripheral blood of mice treated as indicated; n = 3. \**p* < 0.05, \*\**p* < 0.01.

### 3.6 CXCL1^exo-dead^ transcriptionally increases PD-L1 expression in macrophages by activating EED signaling

Next, we sought to investigate the molecular mechanism by which CXCL1^exo-dead^ elevated PD-L1 expression in macrophages. Similar to the function of recombinant murine CXCL1, exo-dead treatment significantly increased the mRNA level of *PD-L1* in Raw264.7 macrophages (**Figure 6A**). Additionally, both recombinant murine CXCL1 and exo-dead could concentration-dependently activate the *PD-L1* promoter activity in Raw264.7 macrophages, which was partially inhibited by CXCL1 NA administration (**Figure 6B**). These results indicated that CXCL1^exo-dead^ transcriptionally induced PD-L1 expression in Raw264.7 macrophages. To characterize the regulators involved in CXCL1-induced *PD-L1* transcription activation, DNA-pull down-MS assay was conducted to detect the deferentially expressed proteins (DEPs) binding with the *PD-L1* promoter region of Raw264.7 macrophages following CXCL1 treatment. The pull-down assay was conducted using the biotin-labelled *PD-L1* promoter fragment while the *PD-L1* promoter-interacting proteins were identified by mass spectrometry. A total of 34 DEPs were identified after CXCL1 treatment. Additionally, the potential transcription factors (TFs) of *PD-L1* gene were predicted using the hTFtarget database. By intersecting the DEP set and the TF set, embryonic ectoderm development protein (EED) was finally determined (**Figure 6C**). Next, the combined effect of CXCL1 and EED knockdown on PD-L1 expression in macrophages was further investigated to validate whether EED was the potential transcription factor responsible for CXCL1-induced *PD-L1* transcription. CXCL1 significantly induced the expression and nuclear translocation of EED, and therefore activate the promoter activity and protein expression of PD-L1 in Raw264.7 macrophages (**Figure 6D–F**). However, EED knockdown partially abrogated CXCL1-induced PD-L1 transcription and protein expression (**Figure 6E–F**). Next, the binding sites of EED in the *PD-L1* promoter region as well as the binding activity changes after CXCL1 treatment were further investigated to better elucidate the molecular mechanism. JASPAR prediction suggested that there was one potential EED binding site (5’-GTTCCACTC-3’, –437 to –429 bp) in the *PD-L1* promoter region. CHIP-PCR assay further suggested that CXCL1 significantly promoted the binding of EED with this promoter fragment, while EED knockdown in macrophages partially abrogated their interaction (**Figure 6G**). More importantly, the antisense mutation of the EED binding region in the *PD-L1* promoter dramatically abrogated the induction effect of CXCL1 on *PD-L1* promoter activity in Raw264.7 macrophages (**Figure 6H**). Taken together, these results indicate that CXCL1^exo-dead^ transcriptionally increases PD-L1 expression in macrophages by activating EED signaling.

**Figure 6.**
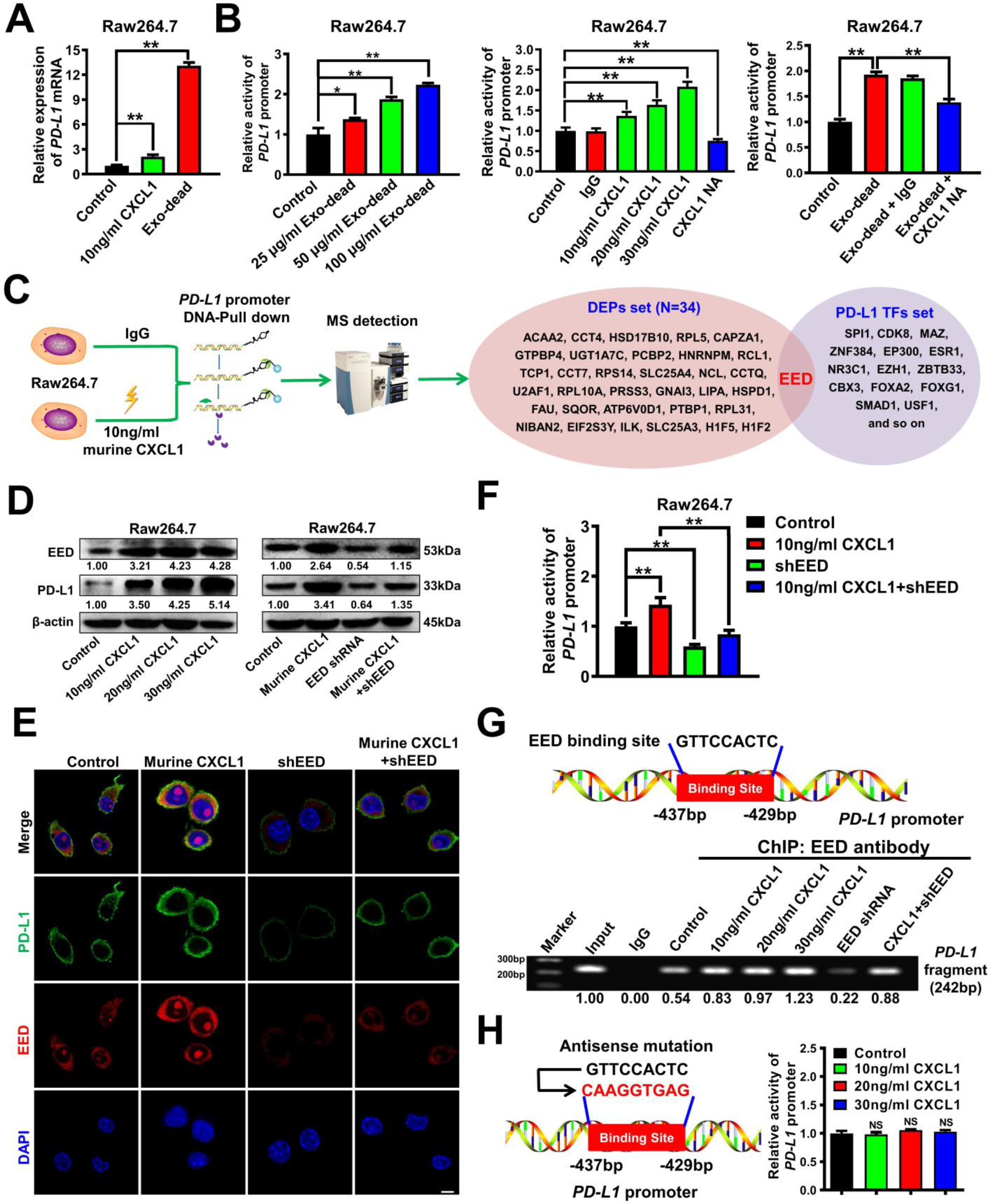
CXCL1^exo-dead^ transcriptionally increases PD-L1 expression in macrophages by activating EED signaling. **(A)** The mRNA level of *PD-L1* in Raw264.7 macrophages was significantly increased by 10 ng/ml murine CXCL1 and 50 μg/ml exo-dead treatment for 24 h; n = 3. **(B)** The promoter activity of *PD-L1* in Raw264.7 cells when treated as indicated for 24 h; CXCL1-NA concentration: 5 μg/ml; n = 3. **(C)** Diagram of the DNA-pull down-MS assay. The DEPs that bind with the *PD-L1* promoter region of Raw264.7 macrophages after CXCL1 treatment for 24 h were analyzed by mass spectrometry. The TFs of *PD-L1* were predicted using the hTFtarget database. EED was selected as the potential transcription factor by taking the intersection of the DEP set and the TF set. **(D–E)** The expression levels of EED and PD-L1 in Raw264.7 cells when treated as indicated for 48 h. Murine CXCL1 concentration: 10 ng/ml. Scale bar: 5 μm. **(F)** The combinational effect of CXCL1 treatment and EED knockdown on the promoter activity of *PD-L1* in Raw264.7 cells when treated as indicated for 24 h; n = 3. **(G)** The binding activity of EED with the promoter fragment of *PD-L1* in Raw264.7 cells when treated as indicated for 48 h was investigated by CHIP-PCR assay. **(H)** The antisense mutation of the EED binding region in the *PD-L1* promoter significantly abrogated the induction effect of CXCL1 on *PD-L1* promoter activity in Raw264.7 macrophages; n = 3. \**p* < 0.05, \*\**p* < 0.01.

### 3.7 TPCA-1 significantly inhibits CXCL1^exo-dead^-induced chemoresistance and invasion of breast cancer cells co-cultured with macrophages

Next, we explored the translational significance of targeting CXCL1^exo-dead^ signaling. As CXCL1 is a chemokine, the commercialized Chemokine Inhibitor Compound Library (TargetMol, Catalog Number: L7600) containing 80 small molecule compounds was screened by ELISA (**Figure 7A**). The top five compounds with the strongest inhibitory activities on CXCL1 secretion from breast cancer cells were identified as C-DIM12, TPCA-1, plerixafor, fudosteine, and AS-604850. TPCA-1 was selected and subjected to further investigations given that it dose-dependently inhibited CXCL1 secretion in 4T1 cells significantly **(Figure 7-table supplement 1** and **Figure 7B**). At concentrations of 1–4 μM, TPCA-1 exhibited little cytotoxicity effects on 4T1 cells (**Figure 7C**), and did not affect the chemosensitivity of 4T1 cells to paclitaxel (**Figure 7D**). Additionally, TPCA-1 had no significant effect on exo-dead secretion from 4T1 cells (**Figure 7E**) but significantly inhibited CXCL1 levels in both the supernatants of 4T1 cells (**Figure 7F**) and the exo-dead of apoptotic 4T1 cells (**Figure 7G**). More importantly, TPCA-1 remarkably reversed the induction effect of exo-dead on macrophage M2 polarization (**Figure 7H**), which suppressed the exo-dead-induced chemoresistance (**Figure 7I**) and invasion (**Figure 7J**) of breast cancer cells in the co-culture system. Taken together, these results identify TPCA-1 as an inhibitor of exosomal CXCL1 secretion, which can significantly suppress the CXCL1^exo-dead^-induced chemoresistance and invasion of breast cancer cells co-cultured with macrophages.

**Figure 7.**
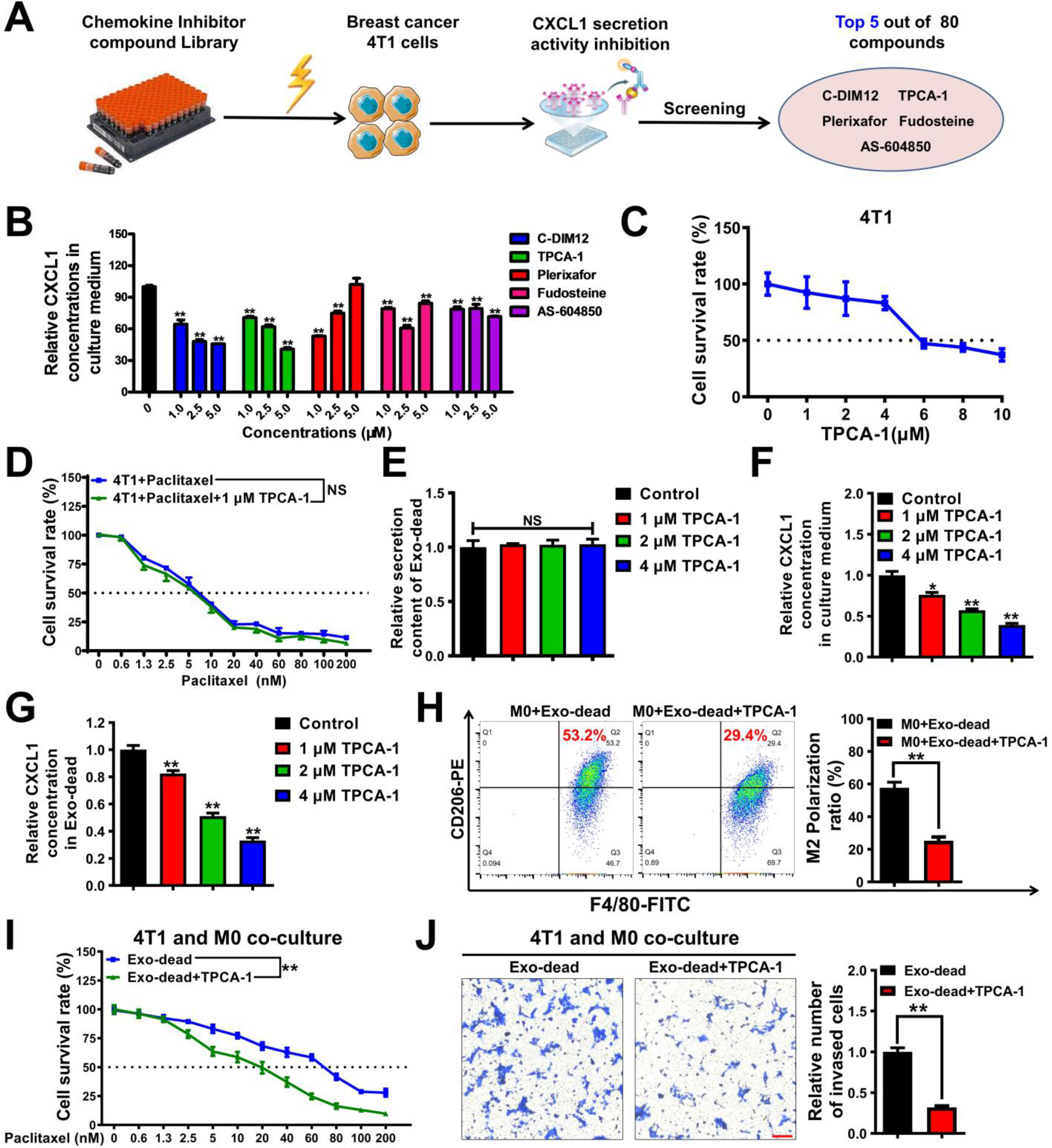
TPCA-1 significantly inhibits exo-dead-induced chemoresistance and invasion of breast cancer cells co-cultured with macrophages. **(A–B)** Diagram of the CXCL1 secretion inhibitor screening assay. 4T1 cells were treated with 80 kinds of compounds (1 μM) for 48 h, and the top five compounds with the strongest inhibitory activities on CXCL1 secretion of 4T1 cells were selected; n = 3. **(C–D)** The cytotoxicity of TPCA-1 in breast cancer 4T1 cells (n = 8) and its effect on the chemosensitivity of 4T1 cells to paclitaxel (n=3). Cells were treated as indicated for 48 h. **(E–G)** TPCA-1 treatment for 48 h had no significant effect on exo-dead secretion (E) from 4T1 cells but significantly attenuated the concentration of CXCL1 in the supernatants of 4T1 cells (F) and the exo-dead of apoptotic 4T1 cells (G); n = 3. **(H)** The results of the flow cytometry assay indicated that 1 μM TPCA-1 treatment for 48 h significantly reversed the induction effect of exo-dead (50 μg/ml) on the M2 polarization of macrophages; n = 3. **(I–J)** CCK-8 and Transwell assays suggested that 1 μM TPCA-1 treatment for 48 h inhibited 50 μg/ml exo-dead-induced chemoresistance (n = 8) and invasion (n = 3) of 4T1 cells in the co-culture system. Scale bar: 100 μm. \**p* < 0.05, \*\**p* < 0.01.

### 3.8 TPCA-1 chemosensitizes breast cancer to paclitaxel and inhibits CXCL1^exo-dead^-induced breast cancer growth and lung metastasis *in vivo*

Finally, the chemosensitizing activity of TPCA-1 was validated *in vivo*. As shown in **Figure 8A–B**, TPCA-1 treatment (10 mg/kg/d) alone moderately inhibited breast cancer growth and lung metastasis in the mouse 4T1-Luc xenograft model. Meanwhile, TPCA-1 treatment significantly inhibited the infiltration and PD-L1 expression of TAMs in the TME **(Figure 8C–D),** and decreased CTCs in the blood of 4T1-Luc xenograft-bearing mice **(Figure 8E)**. More importantly, TPCA-1 could chemosensitize breast cancer to paclitaxel, leading to more significant inhibition effects on breast cancer growth, lung metastasis, TAM infiltration and PD-L1 expression in the TME, and reduced CTC infiltration to the blood **(Figure 8A–E)**. Notably, TPCA-1 treatment at a dose of 10 mg/kg/d for 27 days exhibited no noticeable hepatotoxicity, nephrotoxicity, or hematotoxicity *in vivo* **(Figure 8-table supplement 1**), suggesting the long-term biosafety and the promising druggability of TPCA-1. We also sought to investigate whether TPCA-1 could inhibit the promotion effect of CXCL1^exo-dead^ on breast cancer growth and metastasis. As shown in **Figure 8F–G**, TPCA-1 treatment remarkably suppressed the induction effect of CXCL1^exo-dead^ on breast cancer growth and lung metastasis in mouse 4T1-Luc xenograft, while exo-dead^rCXCL1^ (CXCL1 overloading in exo-dead) blocked the inhibition effect of TPCA-1. Additionally, TPCA-1 treatment also significantly inhibited the promotion effects of CXCL1^exo-dead^ on TAM infiltration and PD-L1 expression in the TME, and CTC infiltration to the blood. However, exo-dead^rCXCL1^ partially abrogated these effects, indicating that the pharmacological effect of TPCA-1 was mainly attributed to CXCL1 inhibition (**Figure 8H–J**). Taken together, these results indicate that TPCA-1 not only chemosensitizes breast cancer to paclitaxel but also inhibits CXCL1^exo-dead^-induced breast cancer growth and lung metastasis *in vivo*.

**Figure 8.**
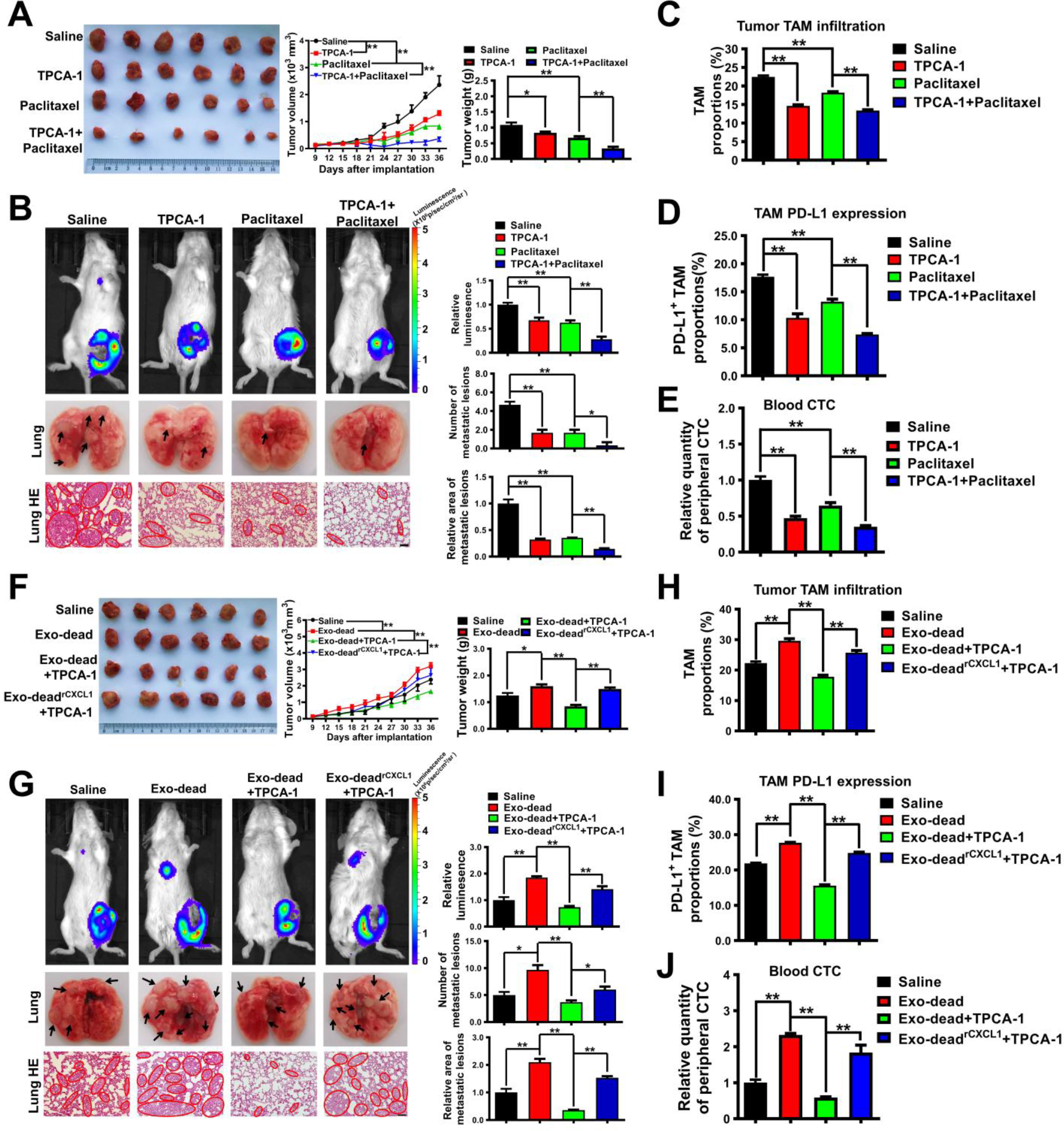
TPCA-1 chemosensitizes breast cancer to paclitaxel and inhibits CXCL1^exo-dead^-induced breast cancer growth and lung metastasis *in vivo*. **(A)** Representative images of tumors and the tumor volume curves (n = 6). TPCA-1 (10 mg/kg/d) and paclitaxel (10 mg/kg/3d) were administered. **(B)** Representative images of the *in vivo* imaging assay and lung HE staining assay; n = 3. Scale bar: 100 μm. **(C–D)** The infiltration levels of CD45^+^/F4/80^+^/CD206^+^ TAMs and CD45^+^/F4/80^+^/PD-L1^+^ TAMs in the TME of mice following treatment with TPCA-1, paclitaxel, or the combination of TPCA-1 and paclitaxel; n = 3. **(E)** The CTC quantity in the peripheral blood of mice when they were treated as indicated; n = 3. **(F)** Representative images of tumors and the tumor volume curves (n = 6). Exo-dead and exo-dead^rCXCL1^ (200 μg/20 g weight, q3d) were administered by peritumoral injection. **(G)** Representative images of the *in vivo* imaging assay and lung HE staining assay; n = 3. Scale bar: 100 μm. **(H–I)** The infiltration levels of CD45^+^/F4/80^+^/CD206^+^ TAMs (n = 3) and CD45^+^/F4/80^+^/PD-L1^+^ TAMs (n = 6) in the TME of mice when they were treated as indicated. **(J)** The CTC quantity in the peripheral blood of mice when they were treated as indicated; n = 3. \**p* < 0.05, \*\**p* < 0.01.

## 4. Discussion

Although chemotherapy represents a cornerstone for breast cancer treatment, emerging evidence has indicated that chemotherapy also plays a key role in mediating cancer metastasis ^7–10^. Indeed, chemotherapy has been reported to increase the infiltration of neutrophils in pancreatic cancer and results in metastasis *via* Gas6/AXL signaling ^39^. Meanwhile, neoadjuvant chemotherapy has been shown to induce breast cancer metastasis by modulating the TME ^7^. Paclitaxel has also been shown to increase CTCs in breast cancer and facilitate metastatic cell seeding in the lung by upregulating the stress-inducible gene *ATF3* in nonmalignant host cells ^40^. Taken together, these findings suggest that a better understanding of chemotherapy-induced metastasis will assist with the development of novel therapeutic strategies to improve cancer prognosis. In the current study, we demonstrate that paclitaxel could promote exosomal CXCL1 signal secretion from dying breast cancer cells, which served to remodel the pro-metastatic TME by polarizing M2 macrophages through activating EED/PD-L1 signaling. More importantly, pharmacological blockage of the exosomal CXCL1 signal in breast cancer cells by TPCA-1 effectively chemosensitized paclitaxel and restrained breast cancer metastasis both *in vitro* and *in vivo* (**Figure 9**). Our findings highlight the important significance of dying cell-released exosomal signals in mediating cancer metastasis. A series of previous studies also demonstrated that dying cell-released components can induce an immunosuppressive TME. Indeed, IL-1α can be rapidly released by necrotic cells to promote malignant cell transformation and proliferation ^41^. Additionally, dying cells contribute to an increase in potassium, which impairs T cell receptor signaling and limits effector T cell responses against cancer ^42^. Meanwhile, emerging evidence has also suggested that exosomes can participate in multiple cellular processes and contribute to cancer development; however, most previous studies have focused on the biological functions of living cancer cell-derived exosomes. Indeed, Wang *et al.* reported that pancreatic cancer-derived exosomal miR-301a could promote pancreatic cancer metastasis by mediating M2 macrophage polarization ^43^. Morrissey *et al.* reported that lung cancer cell-derived exosomes could drive the infiltration of immunosuppressive macrophages within the pre-metastatic niche (PMN) through glycolytic dominant metabolic reprogramming ^44^. The current study demonstrated that exosomes derived from dying breast cancer cells could significantly induce the immune escape and lung metastasis of breast cancer by modulating TAMs, suggesting that dying cell-released exosomes play a crucial role in facilitating the immunosuppressive TME and poor prognosis of cancer. However, whether other types of immune cells are involved in this process needs to be further studied.

**Figure 9.**
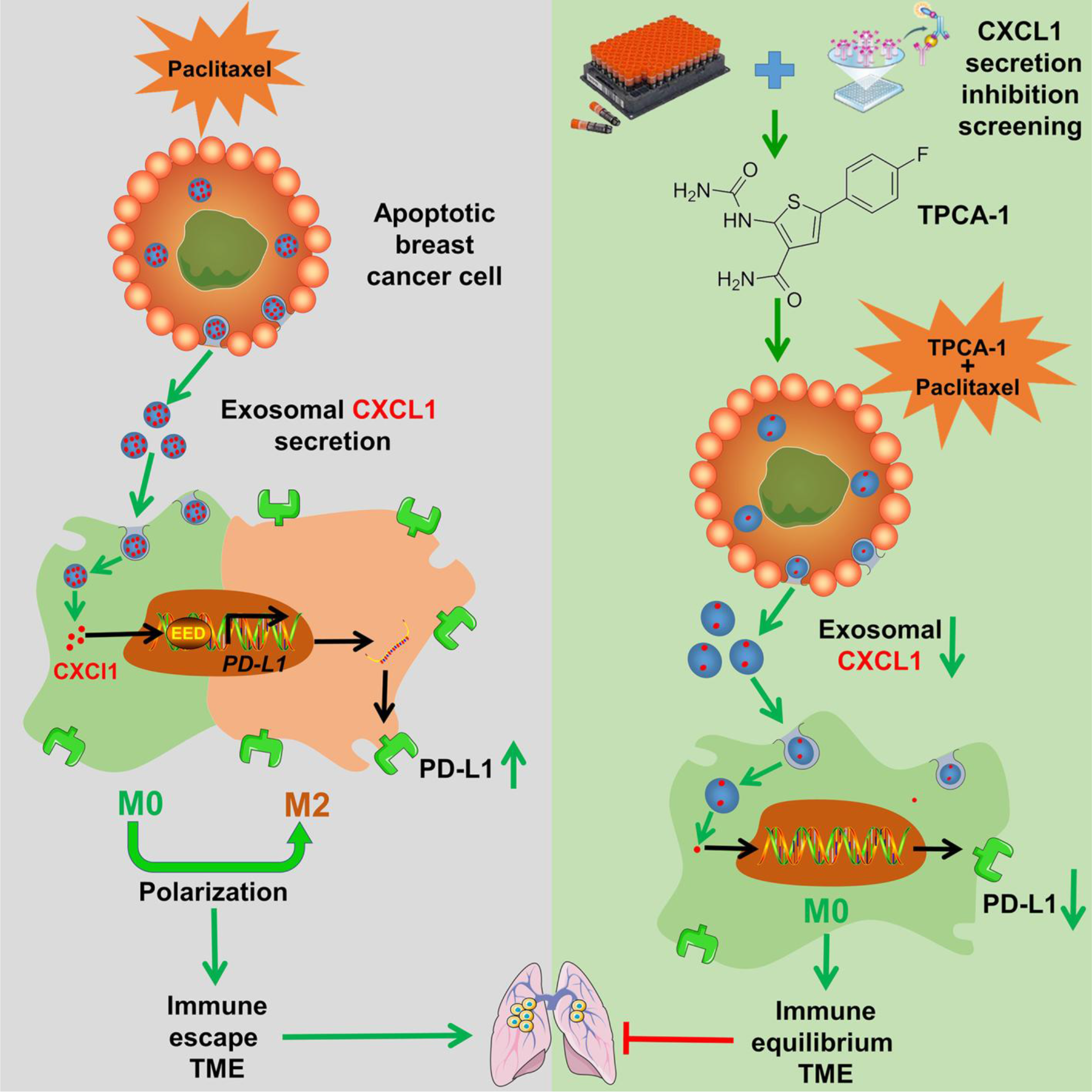
The diagram of the biological process that dying cell-released exosomal CXCL1 signal promotes breast cancer metastasis by activating TAM/PD-L1 signaling as well as its pharmacological blockage by TPCA-1.

In terms of molecular mechanisms, chemokine CXCL1 was found to be enriched in dying cell-released exosomes. CXCL1 represents one of the most abundant chemokines in the TME, and its level in mammary tumor tissue tends to be increased compared to that in normal breast tissue. CXCL1 elevation in breast stroma usually predicts poor OS and recurrence-free survival (RFS) of patients with breast cancer ^36^. CXCL1 can promote breast cancer growth and metastasis through multiple mechanisms, such as inducing epithelial-mesenchymal transformation, promoting the self-renewal of CSCs, inducing autophagy, and accelerating MDSC infiltration and PMN formation ^32^. Our previous study has revealed the important role of TAM-derived CXCL1 in promoting breast cancer metastasis and validated its therapeutic value ^32, 45^. In this study, we demonstrated that paclitaxel could induce the release of exosomal CXCL1 signals from dying breast cancer cells, which is consistent with the previous reports that paclitaxel could increase CXCL1 levels in mice ^46, 47^. However, in contrast to previous studies, we identified that CXCL1 mainly existed in the exosomes. Meanwhile, chemotherapy-induced exosomal CXCL1 signal was identified as the crucial molecular determinant for promoting macrophage M2 polarization and elevating PD-L1 expression to facilitate cancer metastasis. This finding is also consistent with the existing report in Nature showing that chemotherapy-induced necroptosis CXCL1 signals could promote pancreatic oncogenesis and progression by inducing adaptive immune suppression in the TME through activating PD-L1 expression on TAMs ^48^. *PD*-*L1* signaling is an important mechanism utilized by immunosuppressive TAMs to inhibit anticancer responses. Emerging reports have suggested that TAMs represent the major cellular source for maintaining PD-L1 expression in the TME in multiple tumors, including metastatic breast cancer, and that PD-L1 is crucial for the activation and M2 polarization of macrophages ^37, 38^. Molecular elucidation of PD-L1 regulation in TAMs is urgently needed for the successful development of treatment strategies and targeting agents to inhibit breast cancer immune escape. It has been well known that CXCL1 could recruit immune cells and activate their intracellular signal transduction by binding to its receptor CXCR2 ^32, 48^. Therefore, we focused on identifying the downstream molecules that participate in CXCL1-induced PD-L1 expression in TAMs. It was found that CXCL1 induced the expression and nuclear translocation of EED in macrophages, which bound to the 5’-GTTCCACTC-3’ region of the *PD-L1* promoter and transcriptionally elevated *PD-L1* expression. This finding was consistent with the existing reports demonstrating that CXCL1 promoted PD-L1 expression in glioblastoma multiforme cells ^49^ and hepatic cells ^50^ by increasing the activity of the *PD-L1* promoter. Meanwhile, our results also suggest that the combination of PD-L1 blockade and paclitaxel may achieve synergistic inhibition of breast cancer. Indeed, PD-L1 expression in macrophages is associated with the response to neoadjuvant chemotherapy in triple-negative breast cancer ^51^. Meanwhile, PD-L1 expression in residual mammary tumors has been suggested as a prognostic marker in the non-pathological complete response patients after receiving neoadjuvant chemotherapy ^52^. Notably, in the IMpassion 130 clinical study, atezolizumab (anti-PD-L1 antibody) combined with nab-paclitaxel was proven to improve the progression-free survival (PFS) and OS of patients with breast cancer, and this strategy has been clinically approved by the FDA ^53, 54^. Meanwhile, several recently reported neoadjuvant clinical trials incorporating PD-L1 inhibitors with chemotherapy have presented promising results in non-small cell lung cancer ^55, 56^. As CXCL1 was identified as an upstream regulator of PD-L1 in this study, future clinical studies regarding CXCL1 inhibitors plus neoadjuvant chemotherapy are worth investigating in the future.

Based on the above results, we speculated that the selective inhibition of exosomal CXCL1 signaling by small molecule inhibitors may be a promising treatment strategy to enhance chemoresponse and inhibit exosomal CXCL1-induced breast cancer metastasis. In this study, TPCA-1, a selective IκB kinase (IKK) inhibitor, exhibited the strongest inhibitory activity on CXCL1 secretion among 80 compounds from the chemokine inhibitor library. TPCA-1 is a well-known inflammation inhibitor, which functions by inhibiting the release of inflammatory cytokines (e.g., TNF-α, IL-1β, IL-6, and IL-8) and inactivating NF-κB and STAT3 pathways. Notably, TPCA-1 has attracted increasing attention recently, and preclinical studies have proven the therapeutic efficacy of TPCA-1 in autoimmune and inflammatory diseases, such as rheumatoid arthritis, periodontitis, rhinitis, and pneumonia ^57^. However, existing reports on the anti-cancer activity and mechanisms of TPCA-1 have been limited until now. Nan *et al.* reported that TPCA-1 could inhibit mutant EGFR-associated human non-small cell lung cancer by inactivating the STAT3 and NF-κB pathways ^58^. Moreover, using transcriptome-based drug repositioning, TPCA-1 was also screened as a potential selective inhibitor of esophagus squamous carcinoma ^59^. Here, we report for the first time that TPCA-1 could significantly chemosensitize paclitaxel and inhibit CXCL1^exo-dead^-induced growth and metastasis of breast cancer. Our results highlight the development value of TPCA-1 in inhibiting dying-cell-released cytokines during chemotherapy. More importantly, TPCA-1 showed limited hepatotoxicity, nephrotoxicity, and hematotoxicity *in vivo,* suggesting that it may be safely used along with chemotherapy. These results suggest that TPCA-1 may be developed as an adjuvant agent co-administrated with chemotherapy to clean dying cell-released signals and improve prognosis. However, in-depth pre-clinical and clinical studies are still required to validate the value of TPCA-1 in cancer therapy and examine its druggability.

## Conclusion

Taken together, our study demonstrated that dying breast cancer cells induced by paclitaxel could secrete exosomal CXCL1 to promote breast cancer growth and metastasis by activating TAM/PD-L1 signaling. Furthermore, we demonstrated that TPCA-1 could inhibit CXCL1^exo-dead^ signals and chemosensitize breast cancer to improve prognosis. Our findings not only delineate the novel biological mechanism of CXCL1^exo-dead^/TAM/PD-L1 signaling in dying cell-induced immunosuppressive TME but also highlight the potential use of TPCA-1 as an exosomal CXCL1 inhibitor to chemosensitize breast cancer and limit dying cell signal-induced metastasis.

## Acknowledgments

This study was supported by the National Natural Science Foundation of China [82074165, 82174165, 81873306]; the Natural Science Foundation of Guangdong Province [2022A1515011412]; the State Key Laboratory of Dampness Syndrome of Chinese Medicine [SZ2021ZZ19]; the 2020 Guangdong Provincial Science and Technology Innovation Strategy Special Fund (Guangdong-Hong Kong-Macau Joint Lab) [2020B1212030006]; the Science and Technology Planning Project of Guangdong Province [2017B030314166]; Guangdong Science and Technology Department [2021A0505030059]; Guangzhou Science and Technology Project [202201020357, 202102010316]; the Young Elite Scientists Sponsorship Program by CACM [2021-QNRC2-B02], the Research Fund for Qingmiao Talents of Guangdong Provincial Hospital of Chinese Medicine [SZ2022QN01] and the Research Fund for Bajian Talents of Guangdong Provincial Hospital of Chinese Medicine [BJ2022KY12].

## Author contributions

ZYW and SQW supervised the project and wrote the manuscript. SQW and JL performed the experiments and analyzed the data. NW, BWY, YFZ, XW, JPZ and BP took part in the discussion and proofreading of the manuscript. All authors read and approved the final manuscript.

## Data availability

No datasets were newly created or reused in this study. All data generated or analysed during this study are included in the manuscript and supporting file; Source Data files have been provided for all figures and tables (Figures 1-8, figure supplement 1 and table supplements 1-2).

## Supplementary files

**Figure 1-figure supplementary 1.** Effects of exo-alive and exo-dead on the proliferation of breast cancer 4T1 cells. The viability of 4T1 cells after exo-alive and exo-dead treatment for 48 h was investigated using CCK-8 assay *in vitro* (n = 8). The effects of exo-alive (100 μg/ml) and exo-dead (100 μg/ml) on the proliferation and metastasis of 4T1 cells in zebrafish (n = 6).

**Figure 7-table supplement 1. The inhibitory effects of 80 kinds of small molecules on CXCL1 secretion from 4T1 cells.**

**Figure 8-table supplement 1. TPCA-1exhibited no noticeable hepatotoxicity, nephrotoxicity, or hematotoxicity *in vivo*.**

## Source data

**Source data 1. Figures 1-8 source data.**

Figure 1-source data 1. The source data related to Figure 1.

Figure 1-source data 2. Uncropped and labelled blots from Figure 1A.

Figure 2-source data 1. The source data related to Figure 2.

Figure 3-source data 1. The source data related to Figure 3.

Figure 3-source data 2. Uncropped and labelled blots from Figure 3D-E.

Figure 4-source data 1. The source data related to Figure 4.

Figure 4-source data 2. Uncropped and labelled blots from Figure 4A.

Figure 5-source data 1. The source data related to Figure 5.

Figure 5-source data 2. Uncropped and labelled blots from Figure 5A.

Figure 6-source data 1. The source data related to Figure 6.

Figure 6-source data 2. Uncropped and labelled blots from Figure 6D.

Figure 6-source data 3. Uncropped and labelled gels from Figure 6G.

Figure 7-source data 1. The source data related to Figure 7.

Figure 8-source data 1. The source data related to Figure 8A-E.

Figure 8-source data 2. The source data related to Figure 8F-J.

**Source data 2. Figure supplement source data.**

Figure 1-figure supplementary 1-source data 1. The source data related to Figure 1-figure supplementary 1.

Figure 7-table supplement 1-source data 1. The source data related to Figure 7-table supplement 1.

Figure 8-table supplement 1-source data 1. The source data related to Figure 8-table supplement 1.

**Source data 3. Original images of gels and blots.**

